# The outer surface protease, SepM, is required for *blp* locus activation in three of the four most common pherotypes of *Streptococcus pneumoniae*

**DOI:** 10.1101/2022.05.27.493805

**Authors:** Samantha Ratner, Kevin Bollinger, John Richardson, Suzanne Dawid

## Abstract

*Streptococcus pneumoniae* (pneumococcus) is an important human pathogen that primarily resides in the nasopharynx. To persist in this polymicrobial environment, pneumococcus must compete with other members of the bacterial community. Competition is mediated in part by the action of the *blp* locus which encodes a variable array of bacteriocins and their associated immunity proteins. The locus is controlled by a two-component regulatory system that senses the extracellular concentration of the peptide pheromone, BlpC. There are four major pherotypes of BlpC that can be found in most pneumococcal genomes. Here, we show that the protease, SepM, is required for activation of three of the four major pherotypes. The only SepM independent BlpC type is 9AA shorter than the SepM-dependent peptides, consistent with a cleavage event at the C-terminal end. The processing event occurs following secretion and removal of the C terminal region is required for binding to the histidine kinase receptor. Synthetic truncated peptides or full-length peptides pre-incubated with SepM-expressing bacteria can upregulate the *blp* locus independent of SepM. We show that SepM-independent peptides accumulate in the supernatant of secreting cells at low levels suggesting a role for the tail in peptide secretion, stability or solubility and demonstrating a significant tradeoff for SepM-independence.

**Importance:** *Streptococcus pneumoniae* is an important cause of disease in humans that occurs when the bacteria in the nasopharynx bypasses host defenses to invade deeper tissues. Colonization fitness thus represents an important initial step in pathogenesis. *S. pneumoniae* produces antimicrobial peptides called bacteriocins which provide a competitive advantage over neighboring bacteria in the nasopharynx. The *blp* locus encodes a variable array of bacteriocins that participate in competition. Here, we demonstrate that activation of the *blp* locus requires a surface protease that activates the *blp* signal peptide. There are naturally occurring signal peptides that do not require cleavage, but these are characterized by poor secretion. We describe an additional, previously unappreciated activation step in the control of bacteriocin production in *S. pneumoniae*.

## Introduction

*Streptococcus pneumoniae* is a human pathogen that resides primarily in the human nasopharynx as a member of the resident microbial community. Breaches in host barriers allow for invasion of deeper tissues and symptomatic disease, but colonization is typically asymptomatic. Young children serve as the major reservoir for this bacterium and are typically serially colonized with different *S. pneumoniae* strains over time ^1–4^. Some individuals are colonized by a single strain, but a single child can be colonized with as many as 5 distinct strains at any given time^5,6^. Genome studies demonstrating evidence of DNA exchange between pneumococci at the individual and population level confirm that pneumococcus regularly encounters and competes with other pneumococci the nasopharynx^7,8^. Competition with other members of community is likely to also play an important role in the ability of an isolate to establish itself on the mucosal surface. To compete, *S. pneumoniae* produces antimicrobial peptides called bacteriocins. The *blp* locus is found in all pneumococcal genomes and encodes a variety of single and two peptide bacteriocins called pneumocins that target other pneumococci and closely related streptococci^9–14^. *blp* activity is controlled by a two-component regulatory system (BlpRH) that is activated by the detection of a peptide pheromone called BlpC^11,15^. In the prototypic strain, BlpC is secreted out of the cell by the PCAT family transporter complex, BlpAB. The leader peptide of pre-BlpC, up to a conserved double glycine motif, is cleaved by the protease domain of BlpA upon transport out of the cell. BlpC binds its cognate receptor, the histidine kinase, BlpH which results in a conformational change and dimerization. A resulting phosphotransfer event activates the response regulator, BlpR. The phosphorylated regulator binds to inverted repeats within and outside of the *blp* locus, upregulating the transcription of the operons encoding BlpAB and BlpC in addition to the region encoding the pneumocins and their associated immunity proteins referred to as the Bacteriocin Immunity Region (BIR). Like BlpC, pneumocins are characterized by a typical leader sequence followed by a double glycine motif and in the prototypic strain are processed and secreted out of the cell by the BlpAB transporter complex.

The *blp* locus is variable from genome to genome suggesting that selective pressure is shaping the bacteriocin/immunity content in addition to aspects of locus regulation^12–14,16,17^. The *blp* locus itself has been shown to represent a hot spot of recombination within co-colonized individuals^8^. The BIR of individual strains can encode up to six of the ten described distinct bacteriocin peptides with their co-transcribed immunity proteins. Allelic variants of bacteriocin genes have been shown to impact inhibitory activity^9,10^. The ORF encoding the major transporter subunit, BlpA is also characterized by variability. Depending on the genome collection assessed, nearly 50% of genomes carry a frame shifting 4 bp repeated sequence that results in a nonfunctional transporter. An additional 20-25% of strains have a large deletion of the *blpA* gene or other inactivating mutations leaving approximately a quarter of isolates with an intact and functional BlpAB transporter^14,16,18^. Despite the absence of a dedicated transporter, BlpA deficient strains can use the homologous competence regulated ComAB transporter to secrete both BlpC and the pneumocins. The ComAB transporter is produced at low basal levels and is briefly upregulated during the development of competence. Competence in *S. pneumoniae* is controlled by the ComDE two-component system that is activated by accumulation of the peptide pheromone CSP. CSP is secreted out of the cell by ComAB where it binds to the ComD histidine kinase^19–21^. Like the BlpRH system, ComD dimerization results in a phosphotranfer reaction that ultimately activates the response regulator, ComE. Phosphor-ComE upregulates the transcription of several early competence genes including the *comAB* and *blpABC* operons^22,23^. In the 75% of strains that lack a functional BlpAB complex, *blp* activation always follows a peak in competence gene activation^24^. In the strains that express a functional BlpAB complex, *blp* activation can occur separately from competence, and because BlpAB can also contribute to the secretion of CSP, competence occurs at lower cell densities and under less permissive conditions. In addition, these strains secrete significantly more pneumocins over time and show a clear pneumocin-mediated competitive advantage in plate antagonism assays and during co-colonization of biofilms and in colonizing the mouse nasopharynx^25^. In surveying available genomes, *blp* loci also differ by their *blpC/blplH* alleles. There are four major BlpC types that are paired with specific BlpH types^17,26^. BlpC binding to BlpH has been shown to involve the association of the N-terminal region of the peptide with the first ∼150AA of BlpH^17^. Different BlpC types do not participate in cross talk with other BlpH types with some exceptions. The restriction of cross talk presumably occurs to limit shared signaling between competing pneumococcal strains.

In this work, we show that three of the four major BlpC types must be processed at the C-terminus for BlpH binding. This processing requires the presence of the conserved outer surface protease, SepM. SepM cleavage is absolutely required for *blp* activation in vitro using full-length synthetic BlpC and for natural activation during growth. Based on the demonstrated SepM specificity in other strep species, we conclude that SepM removes the C-terminal 7 amino acids. One major BlpC type is produced in a naturally shortened form, lacking the 9 C-terminal amino acids found in longer types and does not require SepM activity to activate the *blp* locus. We show that this peptide is found in low concentrations outside of the cell compared with the SepM dependent pheromones, hence, the tradeoff for SepM independence is low efficiency secretion.

## Results

### Pre-incubation of BlpC_R6_ and BlpC_6A_ with cells enhances *blp* signaling

In the setting of a different line of investigation, we noted that preincubation of BlpC types 6A and R6 with a *blp* locus deletion strain resulted in enhanced signaling when supernatants were used to activate BlpC specific reporter strains compared with peptides incubated in media alone (**Error! Reference source not found**.). The BlpC_R6_ and BlpC_6A_ reporters used here carry a deletion in the *blpC* gene to prevent positive feedback upregulation. Because the incubating strain lacked the entire *blp* locus including the gene encoding BlpC but was otherwise an identical background to the reporter strains used, this suggested that the peptide itself was modified during incubation by an enzyme that is not within or regulated by components of the locus. Of the four major BlpC types found encoded in pneumococcal genomes, BlpC_R6_, BlpC_6A_ and BlpC_P164_ are all 27AA long following the GG motif; only BlpC_T4_ is 9AA shorter at 18AA (Table 1). When we preincubated BlpC_T4_ with cells, there was no enhancement in in *blp* activation compared with media incubated peptide. This observation led us to speculate that proteolytic cleavage was responsible for enhancement of the two longer forms.

**Table 1.**
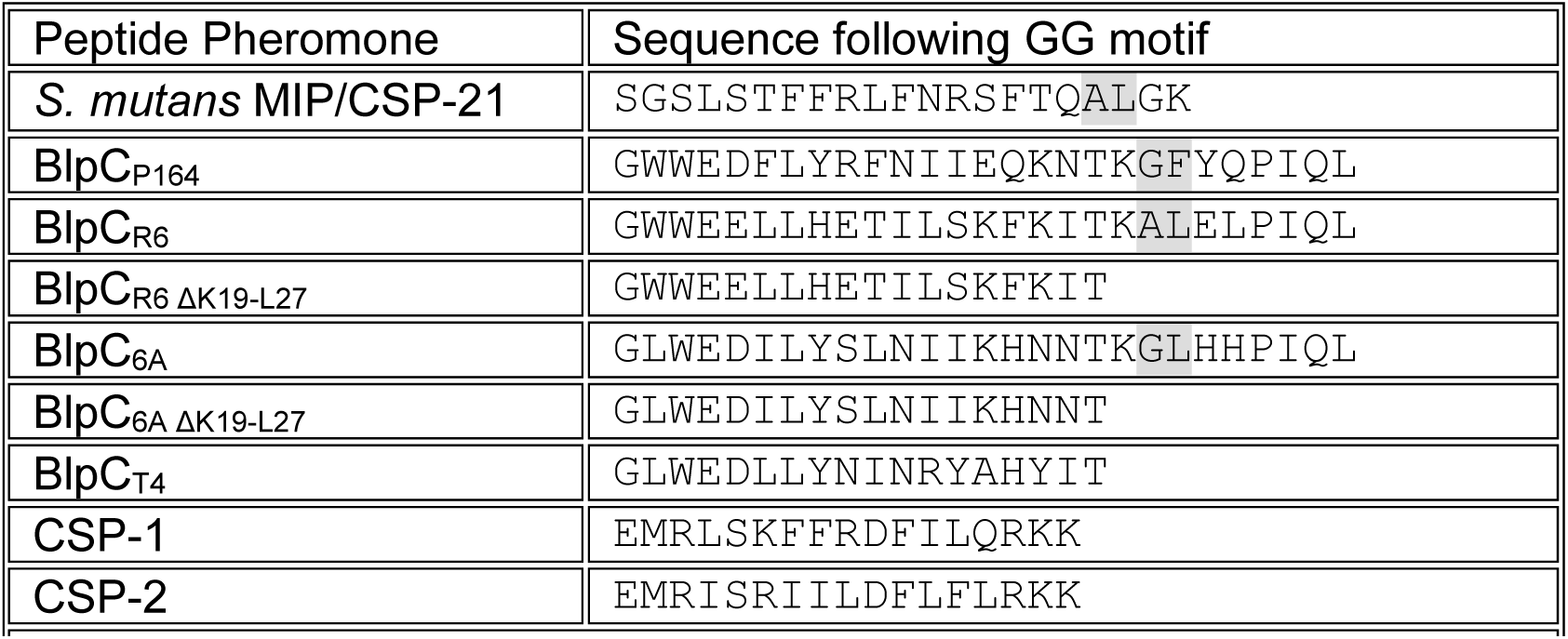
Peptide sequences following cleavage at the GG motif by PCAT proteases. Presumed SepM cleavage sites are highlighted in grey.

### Removal of the C terminal 9AA of BlpC_6A_ and BlpC_R6_ recapitulates pre-incubation activity

To determine if peptide cleavage was responsible for the enhancement of activity of the BlpC_6A_ and BlpC_R6_ peptides following incubation with cells, we used synthetic peptides that were designed to recapitulate the size of the naturally occurring BlpC_T4_ by removing the C-terminal 9AA (Table 1). Truncated BlpC_6A_ and BlpC_R6_ reproduced the enhancement noted following incubation with cells. There was no further increase in activity of these two peptides after incubation with cells (Figure 1). Consistent with increased activity of the truncated peptides, dose-response curves comparing BlpC_R6_ and BlpC_6A_ with their truncated version demonstrated a left shift of the curve for both shorter peptides with a roughly 3-4 fold decrease in EC_50_ values when expression levels were measured 20 minutes after peptide addition (Table 2).

**Table 2.** EC50 values calculated for BlpCR6 and 6A types compared with Δ19-27version.

| BlpC type | BlpC<sub>R6</sub> | BlpC<sub>R6 Δ19-27</sub> | BlpC<sub>6A</sub> | BlpC<sub>6A Δ19-27</sub> |
| --- | --- | --- | --- | --- |
| EC<sub>50</sub> (ng/ml) | 4.027 | 0.8748 | 13.60 | 3.527 |
| 95% CI: EC<sub>50</sub> | 3.060 to 5.294 | 0.6716 to 1.140 | 12.07 to 15.34 | 3.198 to 3.890 |
| R squared | 0.9643 | 0.9775 | 0.9933 | 0.9969 |

**Figure 1.**
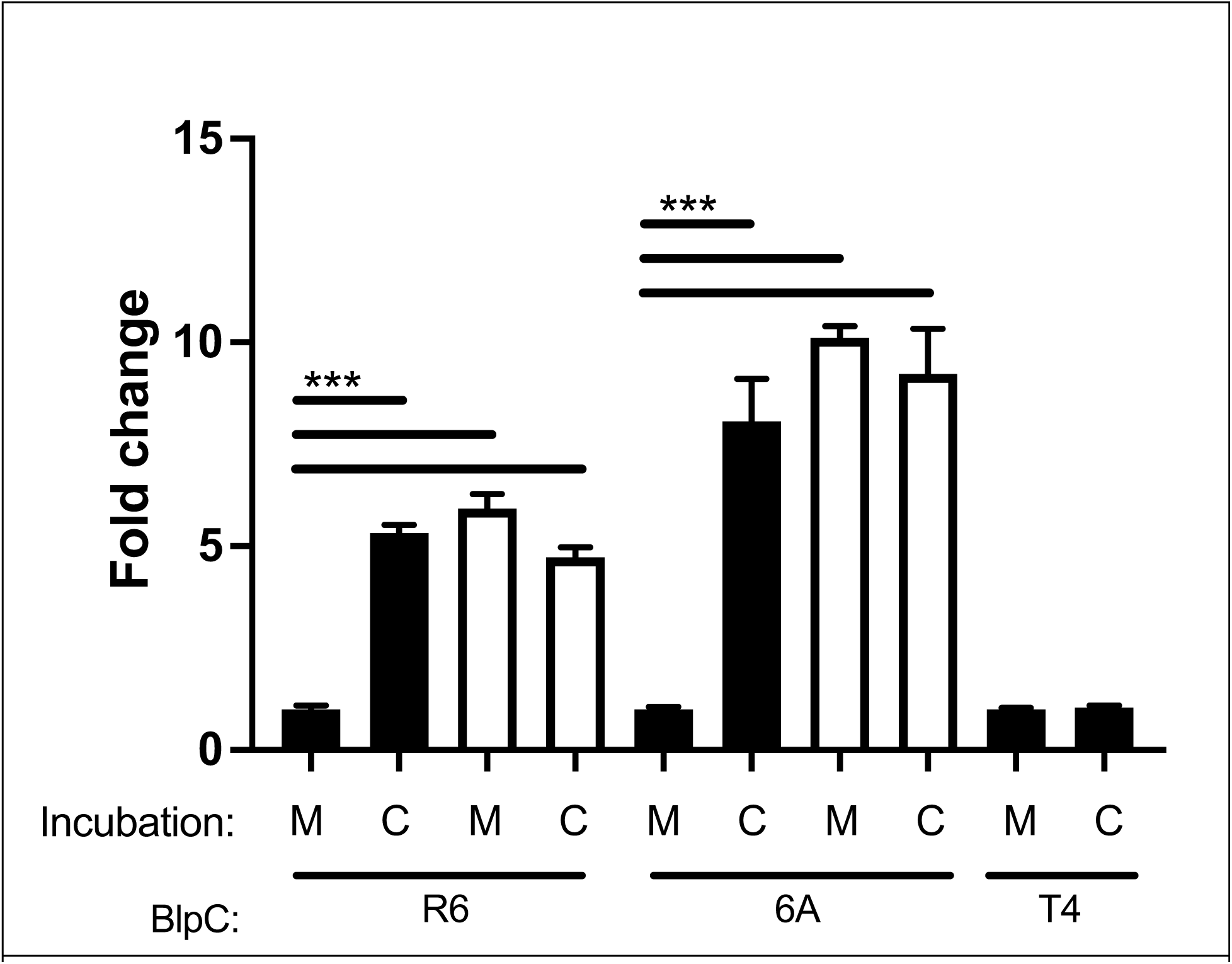
*blp* locus activation with BlpC_R6_, BlpC_6A_ or BlpC_T4_ following pre-incubation of peptide with cells or media. 5 nM of either full length (solid) or Δ 19-27 (open) peptide was incubated with the D39Δ*blpT-X* strain (C) or media alone (M) for 30 minutes at 37C. Cell free supernatants were used to stimulate the type-specific reporter strain and luminescence read at 20 minutes post stimulation. Fold change for each sample compared with full length peptide incubated in media is shown. *** P<.0001 by one way ANOVA.

### *S. pneumoniae* SepM homologue is required for BlpC activation

The CSP-21 peptide pheromone that controls bacteriocin production in *Streptococcus mutans* is cleaved following secretion at its C-terminus by the intramembrane SepM protease to produce the active CSP-18 form^27,28^. The cleavage site is between the A18 and L19 residues, both of which are required for efficient cleavage in this organism. BlpC_R6_ has a similar AL residue at its C terminus. Cleavage at this site would create a peptide just 2AA longer than the tested truncated peptide. BlpC_6A_ and BlpC_P164_ have a GL or GF sequence at this site, respectively. To determine if the *S. pneumoniae* SepM homologue is required for BlpC processing, we tested Δ sepM mutants for *blp* activation. Like in *S. mutans, sepM* is found as the last gene in an operon containing the essential RNA methylase gene *rsmD* and phosphopantetheine adenylyltransferase *coaD* gene^22^. SepM itself is has not been identified as essential. To determine if the SepM homologue is responsible for the activation of BlpC_R6_ and 6A, we created an unmarked deletion of SepM in our reporter strains using allelic exchange of the Janus2 cassette. To address the possibility of unintended mutations, a paired isolate made by retransforming the *sepM::Janus2* strain with the wildtype *sepM* fragment was used as a control. Consistent with a role in activation of full length BlpC_R6_ and BlpC_6A_, Δ*sepM* mutants did not activate following addition of full-length pheromone but response to Δ 19-27 peptides remained intact (Figure 2A, B). Activation of the BlpH_T4_ reporter was unaffected by deletion of *sepM*, consistent with our incubation data suggesting that this form does not require further processing following secretion (Figure 2C). The BlpH_T4_ receptor has been shown to cross stimulate with BlpC_6A_. Consistent with a dependence on SepM for this peptide, only wildtype SepM BlpH_T4_ reporters responded to the addition of BlpC_6A_. No activation was noted in the Δ*sepM* background unless the Δ 19-27 peptide was used (Fig 2C).

**Figure 2.**
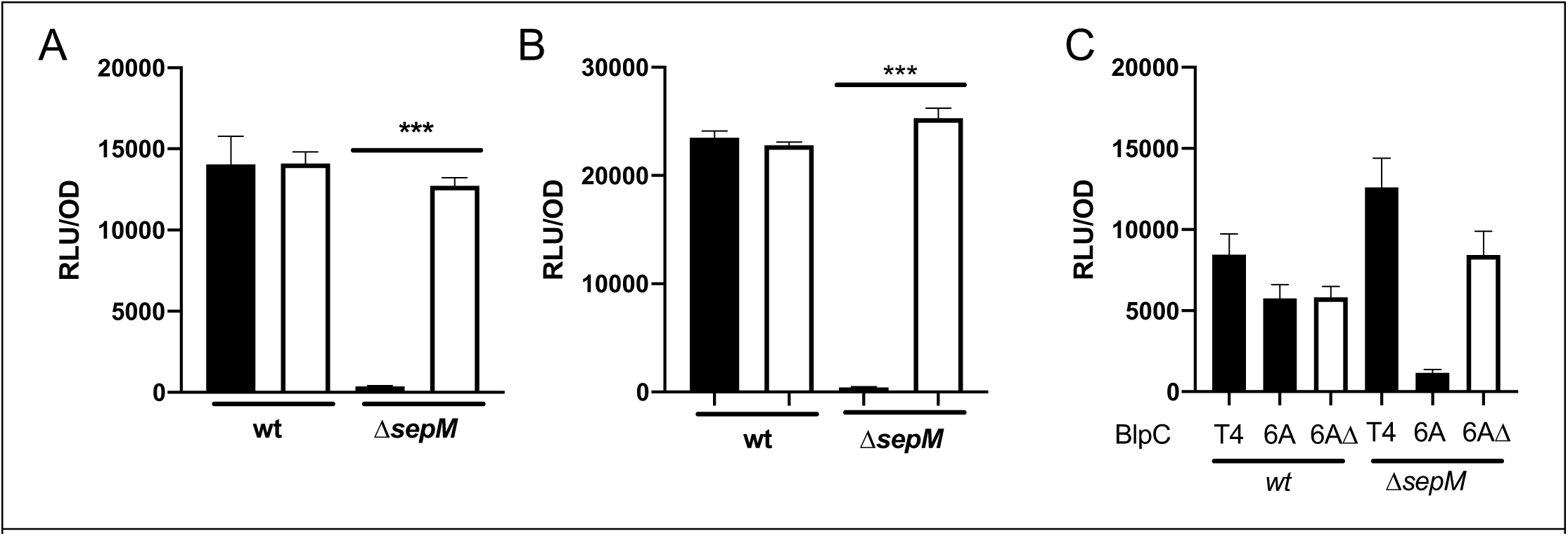
*blp* activation in wt and Δ*sepM* mutant reporter strains in response to synthetic BlpC. pBIR activity one hour after addition of saturating concentrations of either full length (solid) or Δ 19-27 peptide (open) to the peptide-specific Δ*blpC* reporter. A. BlpC_R6_ added to *blpH*_*R6*_Δ*blpC* reporters. B. *BlpC*_*6A*_ added to *blpH*_*6A*_Δ*blpC* reporters. C. BlpC_T4_ or BlpC_6A_ added to *blpH*_*T4*_Δ*blpC* reporters. Luminescence per OD_620_ shown. ***P<.0001 by student T test.

The BlpC_P164_ peptide has the most divergent sequence at the presumed SepM cleavage site (Table 1: GF compared with AL). To determine if activity of this peptide was also dependent on SepM cleavage, we performed a cell incubation assay as in Figure 1 but included a *Δ blpT-XΔ sepM* strain for peptide incubation and a *blpC::spectΔ sepM* BlpH_P164_ reporter strain to detect peptide. Full-length BlpC_P164_ activated the reporter when incubated with the *sepM* wt strain or when the reporter used had an intact *sepM* gene. BlpC_P164_ did not activate the *Δ sepM* reporter strain following incubation with a Δ*sepM* incubating strain, consistent with dependence of this peptide on the activity of SepM for activation (Figure 3). Note that significantly higher, saturating concentrations of BlpC (500 ng/ml) were used in this assay compared with the incubation assays shown in Figure 1 (5 nM = 16 ng/mL) and a later time point was used (20 minutes vs 60 minutes post peptide addition) which accounts for the lack of difference between the wildtype reporter given full length or truncated peptide.

**Figure 3.**
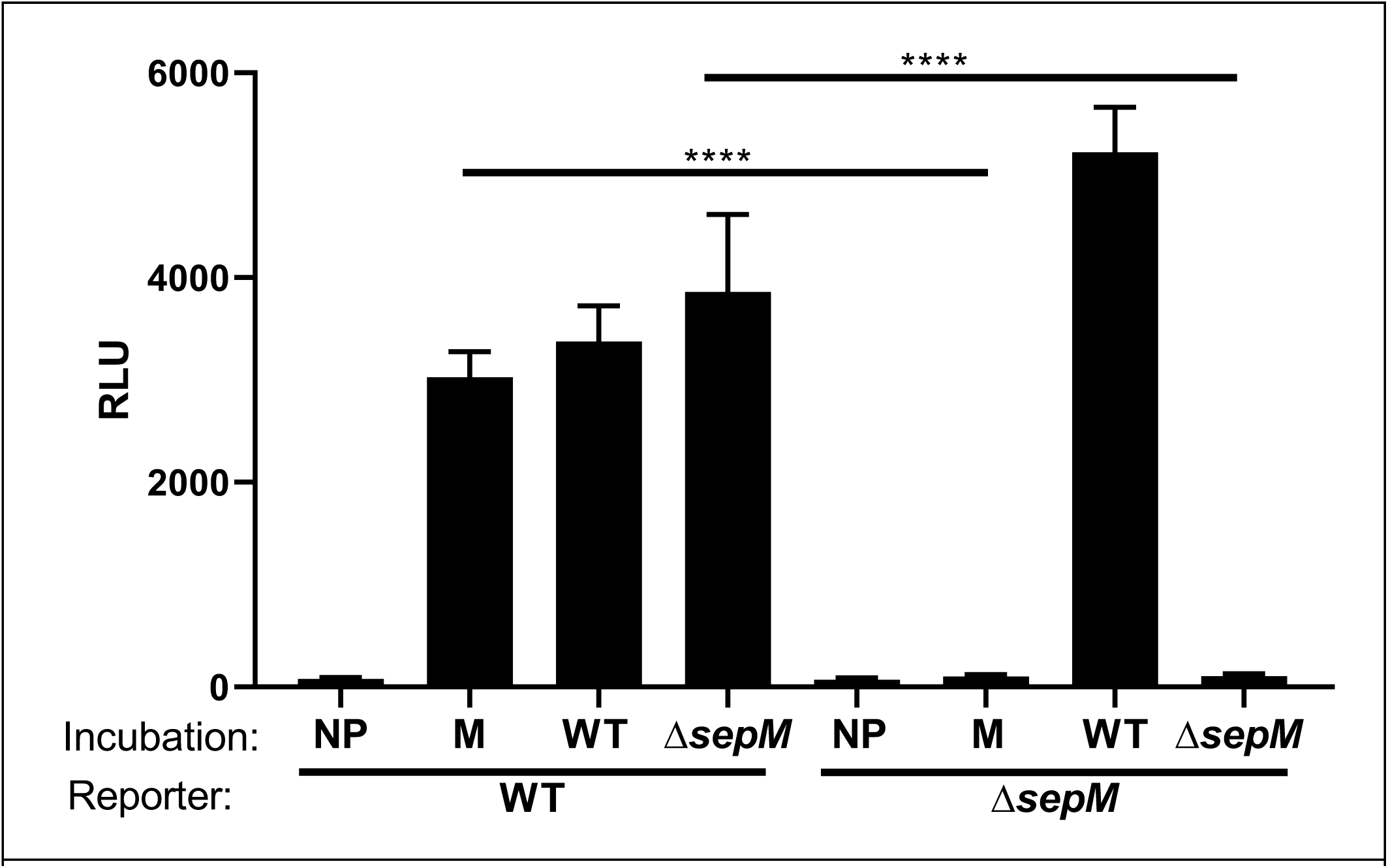
BlpC_P164_ activation of the *blp* locus following incubation with WT, *Δ sepM* cells or media controls. Full length BlpC_P164_ was incubated with a Δ*blpT-X* sepM wildtype strain (WT), a Δ*blpT-X* Δ*sepM* strain, or media alone (M) for 30 minutes. Media without peptide added was used as a baseline control for reporter activity (NP). Supernatants were tested for *blp* activity with the blpH_P164_Δ*blpC* reporter strain carrying either wildtype or *Δ sepM. **** =* P< 0.0001 for the relevant comparisons by one way ANOVA with multiple comparisons.

To determine the impact of a *sepM* deletion on natural *blp* activation, the *sepM* mutation and the replaced wildtype locus was moved into BlpCH_R6_, BlpCH_6A_, BlpCH_P164_ and BlpCH_T4_ backgrounds with intact *blp* signaling systems. Consistent with its role in maturation of the long BlpC pherotypes, we did not appreciate any spontaneous *blp* activation in Δ*sepM* mutants carrying the *blpCH*_*P164*_, *blpCH*_*R6*_ or *blpCH*_*6A*_ alleles (Fig 4 A-C). Reporter strains carrying the *blpCH*_*T4*_ allele demonstrate very small peaks of activation (Fig 4D, E). Consistent with SepM independence, however, there was no difference between the wildtype and *Δ sepM* strains carrying this allele.

**Figure 4.**
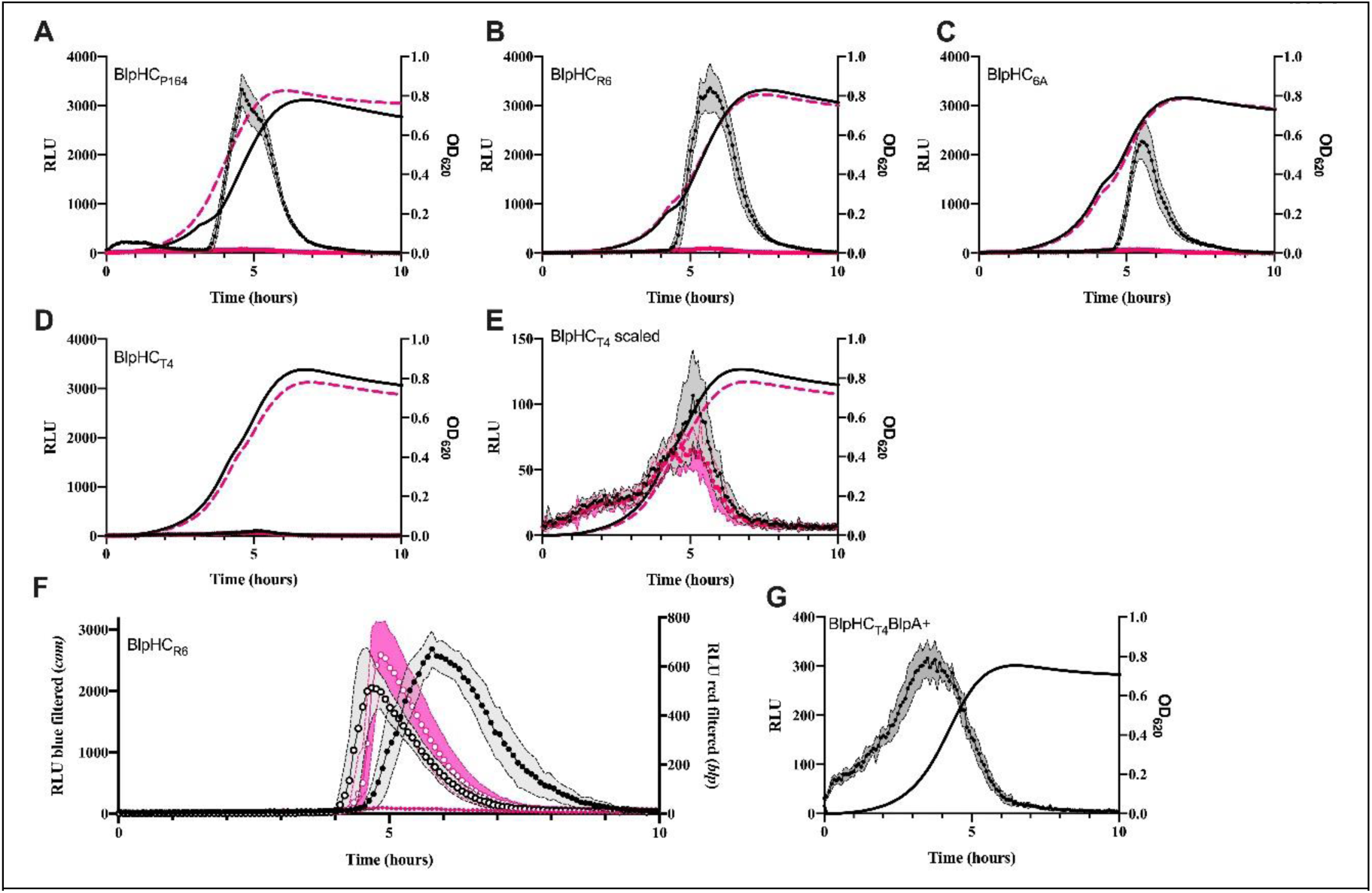
Natural activation of *blp* and *com* reporters with and without SepM. Reporter strains with intact *blp* signaling systems were grown in CDM pH 7.2 media with luciferase substrates. Black circles and lines are wt, pink circles and dashes are Δ*sepM*. Cultures were monitored for growth by OD_620_ (lines) and light production (circles). Points and lines are means and the shaded region encompasses the standard error. In the case of the dual reporter, filters were used to separate the activity of nanolux from firefly luciferase activity. A-D. *blp* reporter strains with the *blpHC*_*P164*_, *blpHC*_*R6*_, *blpHC*_*6A*_ and *blpHC*_*T4*_ alleles were monitored for *blp* locus activation. E. Same data as D but Y axis scaled to show peaks. In F, a dual reporter was used to monitor simultaneously for *com* (open symbols) and *blp* (closed symbols) activation. G. Activation of a *blpA* intact version of the *blpHC*_*T4*_ *blp* reporter strain.

### SepM does not impact CSP mediated competence activation

The competence system in *S. pneumoniae* is controlled by the ComDE two-component regulatory system. CSP is secreted out of the cell by ComAB and BlpAB upon which its leader sequence is cleaved at the GG motif resulting in a 17AA peptide. The two most common CSP pherotypes are CSP1 and 2, both of which have a leucine residue near the C terminus (at -4 or - 5 amino acids from the C-terminus, respectively)^29,30^. Of note, previous work has shown that the C terminal 2-4 AA of CSP1 and 2 are dispensable for their activating activity^30^. If SepM plays a role in competence activation by cleaving CSP, this would impact natural *blp* activation because of the requirement for competence activity in our Δ*blpA* reporter strains. To determine if competence activation requires SepM activity, we assayed for natural activation of competence during growth in competence permissive conditions by using a strain with the *comA* promoter driving nanolux expression with and without the *sepM* gene. Consistent with SepM independence, *comA* promoter activity was identical between wildtype sepM and Δ*sepM* reporter strains. Because these strains are dual *blp/com* reporters the specific effect of the *sepM* deletion on *blp* activation but not competence activation can be clearly seen (Figure 4F).

### The C-terminal extension interferes with binding to BlpH

The inability of full-length BlpC peptides to activate *blp* transcription in a *sepM* mutant could occur because uncleaved BlpC binds to BlpH but fails to create the conformational change required for activation or because the C terminal tail interferes with receptor binding. To address this, we performed competitive dose-response assays with both BlpC_R6_ and BlpC_6A_ full length and truncated peptides using reporter strains carrying the Δ*sepM* deletion. Addition of saturating quantities of full length BlpC_R6_ or BlpC_6A_ did not result in an increase in EC_50_ (Table 3). This lack of antagonistic activity shows that the C terminal extension that is presumably cleaved by SepM interferes with receptor binding. In fact, the EC_50_ values were statistically lower when competing full length peptide was added at saturating conditions. When high concentrations of full length peptide are added to Δ*sepM* strains, there is a small amount of appreciable *blp* activity when compared with reporters given vehicle alone (Supplemental Fig 1). This suggests that there is either a small amount of SepM-independent cleavage or some low level activation activity of intact peptide that can only be appreciated at supra-physiologic concentrations of BlpC.

**Table 3.** EC_50_ values with and without competition using saturating concentrations of full-length BlpC when tested with a Δ*sepM* reporter strain.

| BlpC type | BlpC<sub>R6 Δ19-27</sub> | BlpC<sub>R6 Δ19-27</sub> +<br>BlpC<sub>R6</sub> | BlpC<sub>6A Δ19-27</sub> | BlpC<sub>6A Δ19-27</sub> +<br>BlpC<sub>6A</sub> |
| --- | --- | --- | --- | --- |
| EC<sub>50</sub> (ng/ml) | 1.039 | 0.3683 | 1.448 | 0.9747 |
| EC<sub>50</sub><br>(95% CI) | 0.7174 to 1.507 | 0.2930 to 0.4635 | 1.251 to 1.678 | 0.8260 to 1.151 |
| R squared | 0.9593 | 0.9833 | 0.9933 | 0.9918 |

### Naturally SepM independent BlpC_T4_ is poorly secreted

Based on synthetic peptide assays, BlpC_T4_ is not dependent on the activity of SepM for activation. It was difficult to assess its role in natural activation since our reporter strains with the *blpHC*_*T4*_ alleles show very small peaks of activity during growth in broth culture. To determine if this low activation is related to poor secretion of the peptide, we tested chimeric strains that express mismatched BlpC and BlpH types for BlpC secretion into the supernatant. The secretion activity of a BlpH_R6_BlpC_6A_ chimeric strain was compared with an otherwise isogenic BlpH_R6_BlpC_T4_ chimeric strain. These strains were induced with identical concentrations of synthetic BlpC_R6_ to stimulate *blpC* transcription. After one hour, cell-free culture supernatants were assessed for the concentration of BlpC_6A_ or BlpC_T4_ using their respective *sepM* intact reporter strains. BlpC_R6_ was chosen as the stimulating peptide because it does not stimulate BlpH_6A_ or BlpH_T4_ expressing reporter strains so will not interfere with secreted peptide detection^17^. The concentration of secreted peptide following stimulation with BlpC_R6_ plus or minus CSP was assessed one hour after peptide addition. The activity of filtered supernatants was compared with a dose-response to synthetic full-length peptide. Very little BlpC_T4_ was secreted into supernatants following BlpC_R6_ /CSP stimulation compared with the concentrations secreted by the BlpC_6A_ secreting strain (Fig 5). There is a clear difference in both chimeric strains between the amount of peptide that results from stimulation with BlpC_R6_ alone which would only upregulate the BlpAB transporter and the amount from stimulation with CSP and BlpC_R6_ which would upregulate both BlpAB and ComAB. These data suggest that the tradeoff for SepM independence is a peptide that is poorly secreted into the culture supernatant with barely detectible levels secreted when a single transporter is upregulated.

**Figure 5.**
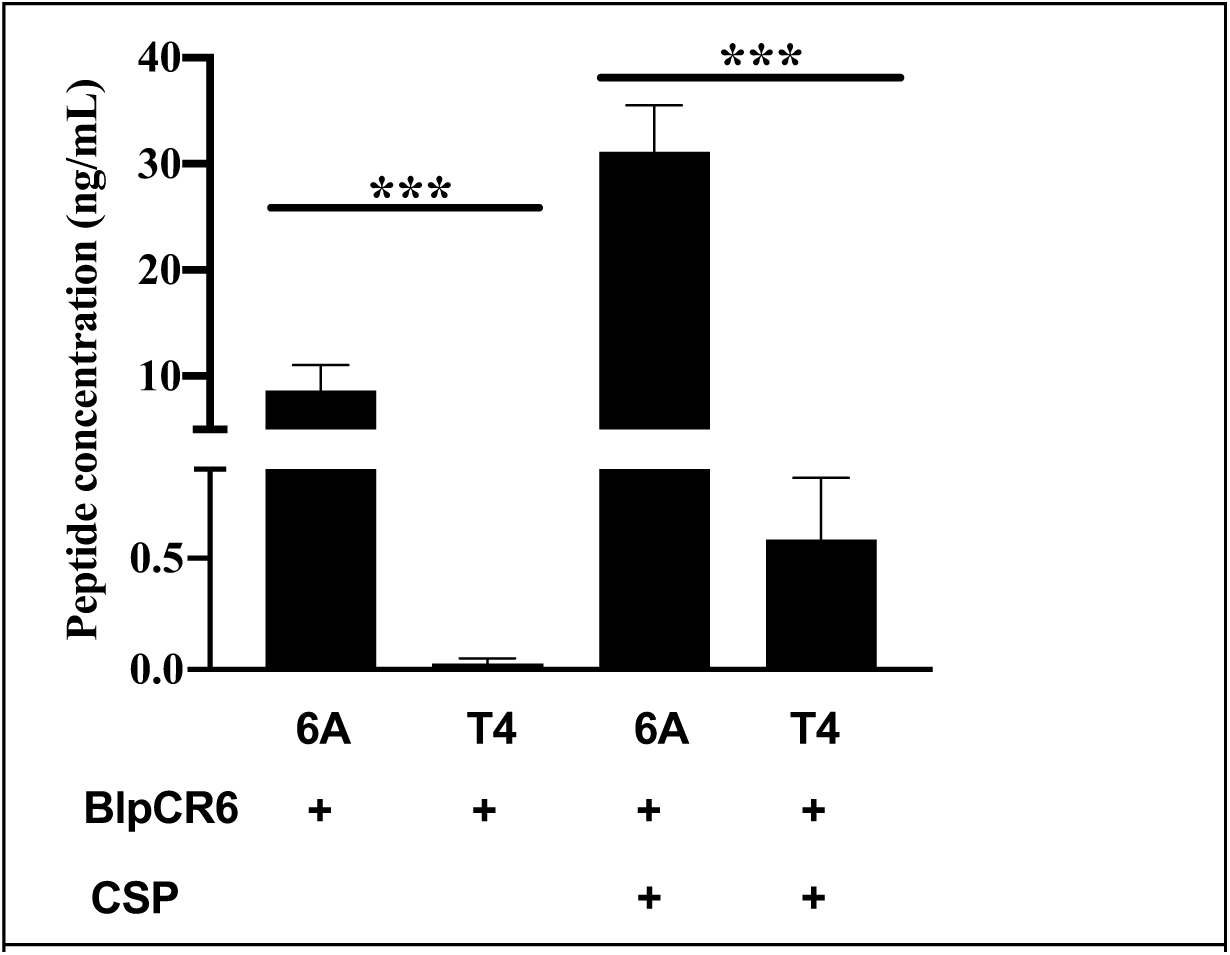
BlpC secretion levels in chimeric BlpC_6A_ and BlpC_T4_ secreting strains. Chimeric strains with either *blpH*_*R6*_*blpC*_*6A*_ or *blpH*_*R6*_*blpC*_*T4*_ were induced with BlpC_R6_ alone or BlpC_R6_ plus CSP. After 1 hour, cells were pelleted and the supernatant was filtered. Cell-free supernatants were assayed for BlpC concentration using pheromone specific reporter strains and concentrations extrapolated from dose-response curve generated with synthetic BlpC. Data analyzed by student T tests. ***P<0.0005

### BlpC_T4_ strains with intact BlpAB show enhanced spontaneous activation in broth culture

The chimera secretion assays demonstrated that very little BlpC_T4_ is secreted when only one PCAT transporter is co-expressed with the peptide pheromone. The reporter strains used to assess for spontaneous induction have the natural frame shift mutation in *blpA*. The BlpC_P164_, BlpC_R6_ and BlpC_6A_ secreting strains can spontaneously activate in this background as long as competence permissive conditions are used. To determine if BlpC_T4_ expressing strains could spontaneously activate at levels closer to the SepM dependent peptides, we used a background with an intact BlpAB. In this background, the *blp* locus activated at substantially higher levels compared with the Δ*blpA* background although peak levels were still lower than noted with the three other BlpH backgrounds (Fig 4G).

### BlpC_6A_ made in a shortened form inside the cell achieves low extracellular concentrations

To better understand the role of the C terminal tail in the SepM dependent BlpC types, we created a version of the BlpH_R6/_BlpC_6A_ chimera with a stop codon engineered in place of L21 to mimic the site of the presumed SepM cleavage event. This peptide would retain signaling activity based on activity of the synthetic truncated peptides but would lack whatever function the C terminal tail plays in intra- and extra-cellular peptide stability, peptide transportation, or peptide solubility. The pre-truncated peptide was found at 1/10^th^ the concentration of the full-length peptide under identical inducing conditions using full length BlpC_6A_ to create a doseresponse for extrapolation (Figure 6). The concentration of the truncated peptide was still quite a bit higher than that seen with the BlpC_T4_ peptide suggesting that other sequence differences are affecting the efficiency of secretion of the T4 peptide.

**Figure 6.**
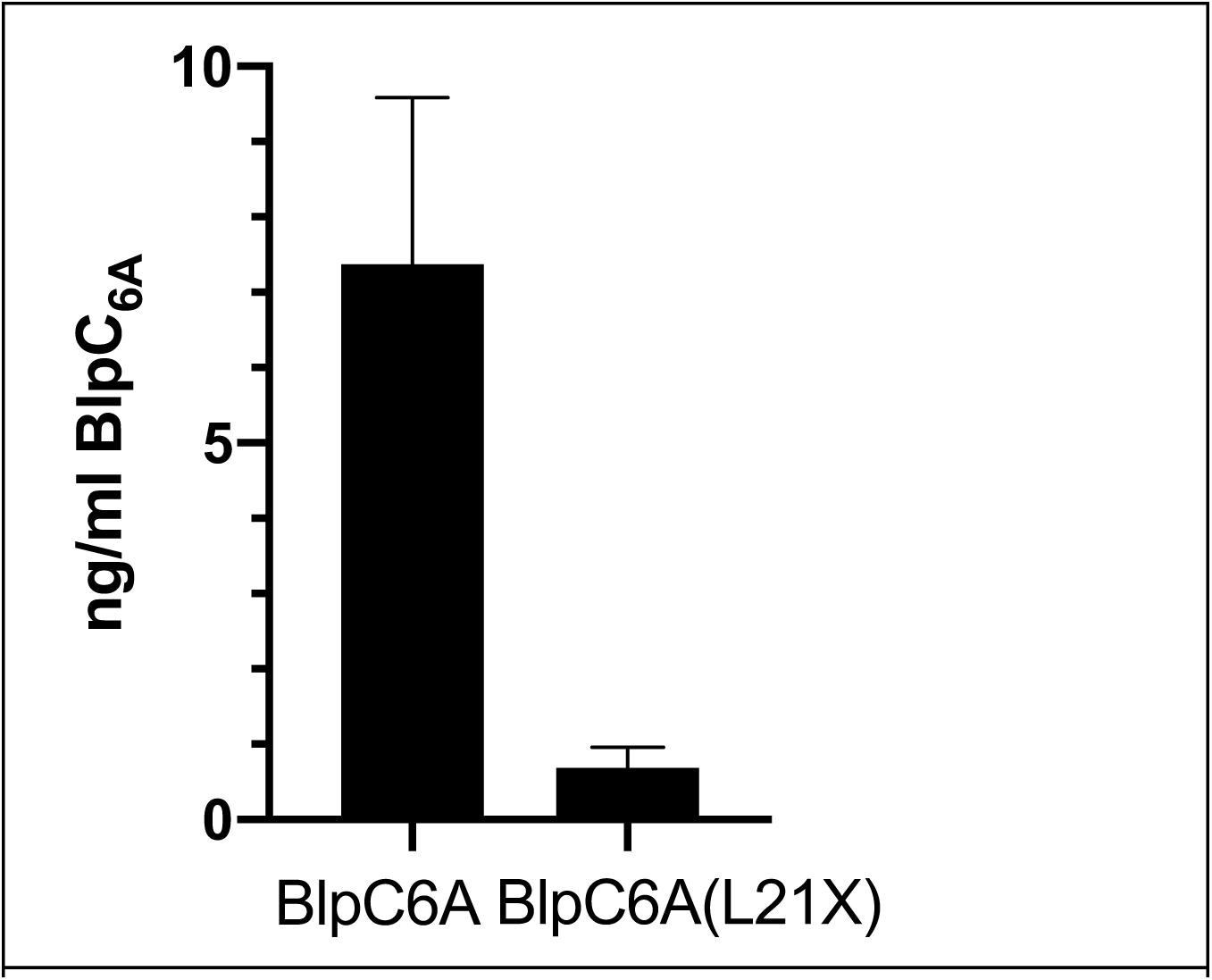
Effect of the removal of the 7 C-terminal amino acids on BlpC secretion. Chimeric strains with either *blpH*_*R6*_*blpC*_*6A*_ or *blpH*_*R6*_*blpC*_*6A133_135CTT>TAA*_ introducing a stop codon at the presumed SepM cleavage site were stimulated with BlpC_R6_ and CSP. After 1 hour, cell free supernatants were tested for BlpC concentration using the BlpC_6A_ specific reporter. Peptide concentrations were extrapolated from a dose-response using synthetic BlpC_6A_. P-value by student T test was < 0.0001

### *S. mutans* UA 159 can process BlpC_R6_ and BlpC_6A_ but not BlpC_P164_

Previous studies on the specificity of *S. mutans* SepM demonstrated a relatively strict requirements at the P1, P’1 cleavage site of SepM substrates to AL^28^. The three 27AA BlpC peptides have AL (R6), GL(6A) and GF(P164) proposed cleavage sites. To determine if *S. mutans* SepM is able to cleave the BlpC peptides into their active form, we incubated saturating concentrations of the three peptides with mid exponential phase growth of either *S. mutans* strain UA159 or our *ΔblpT-X S. pneumoniae* strain or media alone and used cell free supernatants to assay for activation of the type-specific *ΔsepM* reporters (Figure 7). Pre incubated BlpC_P164_ had minimal activity following incubation with *S. mutans*. Higher levels of activity were noted with BlpC_R6_ and BlpC_6A_ following *S. mutans* incubation. These values were all significantly lower than the values obtained following incubation with *S. pneumonia* suggesting that *S. mutans* can process BlpC_6A_ and BlpC_R6_ peptides, but efficiency of processing is lower than seen with the native protease.

**Figure 7.**
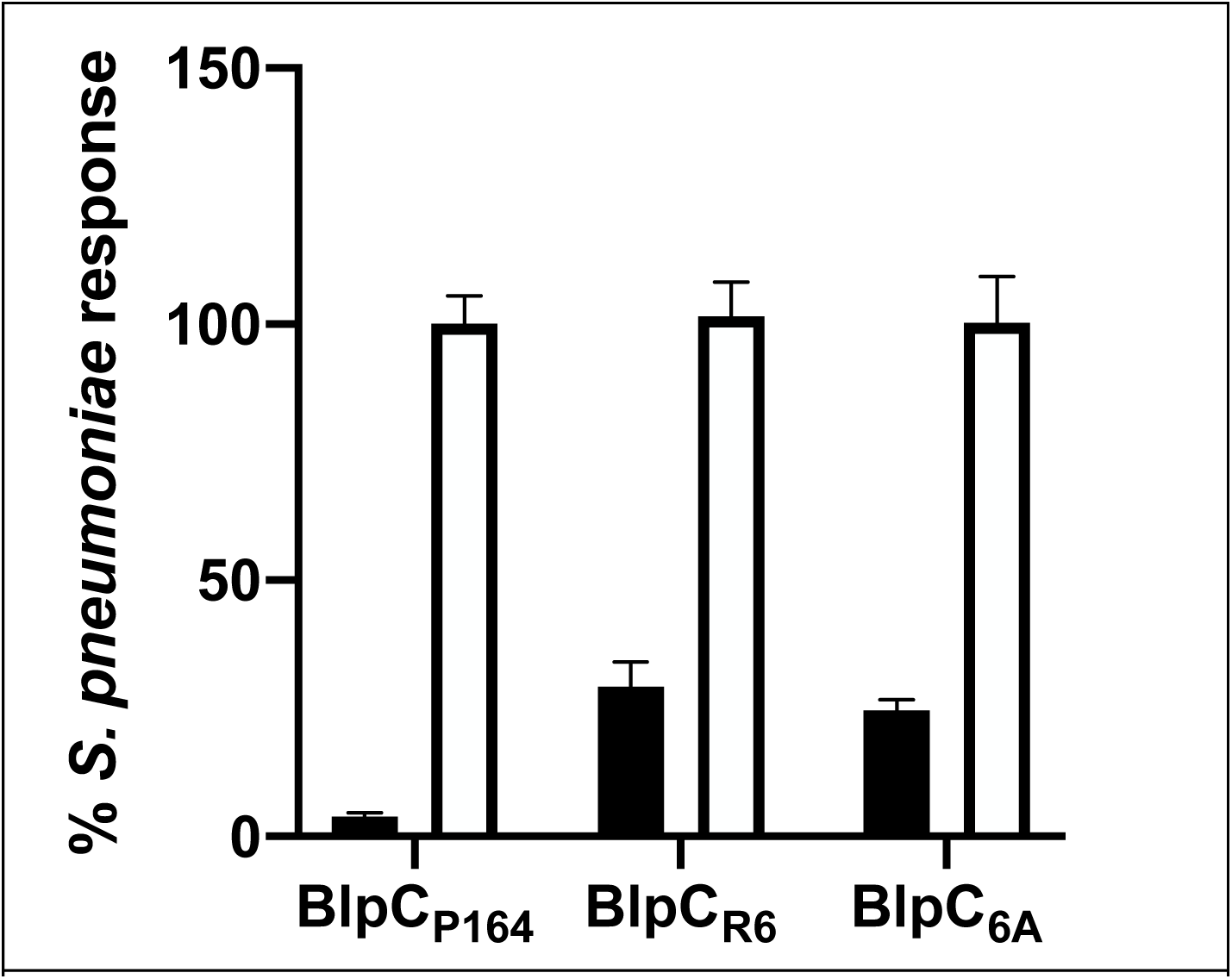
*S. mutans* UA159 incubation assay with the three long forms of BlpC. 500 ng/mL of each of the three 27AA BlpC types were incubated with media alone, *S. mutans* strain UA159 (black) or D39*ΔblpT-X* (white) for 30 minutes at 37C. Cell free supernatants were used to stimulate the Δ*sepM* mutant of the appropriate BlpC reporter strain. RLU values were obtained at 1 hour post stimulation. The media incubated peptide value was subtracted from each value and activation of *S pneumoniae* incubated peptide set at 100%. *S. mutans* incubated peptide values were compared with this value for each peptide/reporter combination. P<0.0001 for each peptide reporter pair by student T test.

## Discussion

SepM is a membrane-localized lon-type serine protease found in most Streptococcal species. As in *S. pneumoniae*, the gene is found as the last gene in a three gene operon in other Streptococcal species^28^. Here, we show that SepM is necessary for *blp* activation in a subset of pneumococcal BlpC pherotypes. Although we did not purify the SepM protease in this study, demonstrating sufficiency for this activation, the most parsimonious explanation of our data is that SepM directly cleaves the longer forms of BlpC. In *Streptococcus mutans*, SepM has been shown to increase the activity of CSP on ComD activation^27^. This *com* locus in *S. mutans* controls bacteriocin production similar to the *blp* locus in *S. pneumoniae*. In S. *mutans*, SepM cleaves the 21AA secreted peptide at the C-terminus removing the last 3 amino acids. Biswas et al. demonstrated that SepM has strict substrate specificity at the P1-P’1 cleavage site where even conserved amino acid changes from AL were found to be inactive^28^. This group specifically tested all amino acids at the P1 site and demonstrated that any change from A was associated with a loss of activation. The group tested three conserved substitutions for the P’1 site (L->V, A or I) and found that none of these were active. Alanine scanning mutagenesis of CSP by Yang et al did not show the same constraints on the P’1 cleavage site sequence but the role of the P1 site was not investigated^30^. Based on this work, we were surprised that the *S. pneumoniae* SepM appears to efficiently cleave at the GF and GL sites found in BlpC_P164_ and BlpC_6A_, respectively. SepM amino acid sequences are only 72% similar and 60% identical when comparing the *S. mutans* and *S. pneumoniae* sequences which may explain the difference in substrate specificity. Consistent with the constraints on cleavage site sequence, we found that preincubation of BlpC_P164_ with *S. mutans* resulted in very small amounts of active peptide whereas BlpC_6A_ and BlpC_R6_ peptides with a leucine residue at P’1 were more efficiently processed. As in *S. pneumoniae, S. mutans* CSP can be found as longer or shorter sequences naturally, but the peptide itself is characterized by less pherotype diversity and thus less variation in the SepM cleavage site in than is found in the pneumococcal *blp* locus.

In *Streptococcus thermophilus*, Boulay et al. demonstrated that SepM is one of three proteases involved in outer surface protein turn over^31^. The authors showed that SepM was either directly or indirectly involved in the cleavage of a number of lipid and membrane anchored proteins. Deletion of SepM in this background did not affect competence or bacteriocin production, but in *S. thermophilus*, both of these systems are controlled by a small hydrophobic peptide (SHP) that is secreted and subsequently imported into the cell where regulator binding occurs. These quorum sensing systems are different from the *blp* system in *S. pneumoniae* and the *com* system in *S. mutans* because BlpC and CSP remain outside of the cell following secretion. In *S. thermophilus*, SepM specificity also favors an alanine residue at the P1 position but is more permissive for the P’1 position. This study examined the whole exopeptidome of *S. thermophilus* with mutations in one or more of the three proteases rather than focusing on cleavage of a single peptide so may have appreciated a wider array of potential cleavage sites because of this. SepM mutants in this background were characterized by abnormal morphology at stationary phase. The authors hypothesize that this phenotype was a result of loss of turnover of proteins involved in cell division suggesting additional roles for this protease in Streptococci. We did not specifically address additional roles of SepM in *S. pneumoniae*, but did note that there were no appreciable growth defects in *Δ sepM* mutant isolates (Figure 4, for example).

We found that SepM is not required for activation of the shorter, BlpC_T4_ peptide. In *S. mutans*, some strains were found to encode a CSP-18 rather than the SepM dependent CSP-21 or 23^28,32^. The authors did not test these strains to determine if the peptide would be efficiently secreted in the absence of the C terminal amino acids. Here, we demonstrate that loss of the C-terminal tail, either in the form of the naturally occurring BlpC_T4_ or in the manufactured early stop codon in BlpC_R6_ significantly impacts extracellular concentrations of peptide. This suggests that the tail-less peptides are less stable inside of or outside of the cell. It is possible that these shorter peptides are degraded or aggregate due to loss of the chaperone-like function of the tail resulting in either poor secretion or inactivity before or after secretion.

BlpC_T4_ is poorly secreted compared with SepM dependent strains but we show it is more efficiently secreted in strains that produce both BlpAB and ComAB compared to those that can only utilize one transporter. BlpC_T4_ encoding strains make up 42% of strains in the Boston isolate collection (a diverse collection of 616 colonizing pneumococcal isolates obtained from children in Boston before and after introduction of Prevnar-7^33^). Interestingly, only 4% of the BlpCH_T4_ encoding strains are predicted to encode a full length BlpA despite an overall frequency of 29% when all *blpHC* alleles are considered. Although our study only addressed the *blpH*_*T4*_ allele from the common strain TIGR4 in the context of our R6 reporter strain, BlpH_T4_ sequences in this collection are characterized by fairly limited diversity, with the majority of isolates having less than 5 amino acids mismatched from the TIGR4 sequence. If our findings are generalizable, this suggests that BlpC_T4_ encoding strains have the specific property of maintaining a very high threshold for activation of the *blp* locus.

The *blp* locus produces bacteriocin peptides that are important in interbacterial competition on mucosal surfaces. The variability in the activity of the locus between isolates as a result of variation in transcriptional activity or pneumocin content suggests that the activity of the locus comes at a significant energetic cost. We have shown that BlpAB-expressing strains demonstrate a competitive advantage in colonization and biofilm models compared with otherwise matched BlpAB disrupted strains^9,10,16,25^. Despite this, these strains only make up a quarter of the population. This suggests that the existing distribution of highly active strains represents an on going balance of competitive gain vs this cost. The requirement for SepM cleavage of BlpC for activation may further contribute to the control of *blp* activation. Based on the *sepM* gene location in an operon with at least one essential gene, it seems unlikely that *sepM* expression is highly regulated. This is supported by pneumoexpress data (https://veeninglab.com/pneumoexpress-app/) showing consistent transcript levels in all growth conditions tested^34^. Protease activity could be impacted at the post-translational level by the presence of competing peptides similar to what is found with the protease, HtrA, whose relative activity in controlling competence is impacted by the concentration of misfolded proteins^35,36^. It is possible that competing substrates may impact relative SepM activity on BlpC further raising the threshold for *blp* activation. This possibility remains theoretical at this point and is an area of active investigation.

## Material and methods

### Growth media and conditions

All *Streptococcus pneumoniae* strains used in this work were derived from D39 or R6 (Table 4). *S. pneumoniae* was grown in THY (Todd Hewitt media supplemented with 5% yeast extract) at pH 6.8 unless otherwise stated. For transformations, cells were grown in C+Y pH 6.8 and stimulated for transformation in C + Y pH 8.0 with 500 ng/mL CSP1 or 2 as appropriate^37^. Either genomic DNA or PCR products were added after 10 minutes incubation at 30C. Outgrowth was carried out prior to plating, first at 30C (40 minutes) then at 37C (1hour). Cells were selected on tryptic soy agar supplemented with 5 μg/ml catalase and antibiotics as follows: gentamicin 200 μg/mL, streptomycin 100 μg/mL, kanamycin 500 μg/mL, spectinomycin 200 μg/mL or media containing 10% sucrose for counter selection of the Janus2 cassette. For natural induction assays, cells were grown in CDM+ pH 7.2. The media contents for CDM+ are described in detail in ^37^. All synthetic peptides were ordered from Genescript (Piscataway, NJ) at >95% purity. Unless otherwise stated, all assays were performed in triplicate on three separate occasions, representative assays are shown.

**Table 4.** Strains used in this study.

| Strain | Properties and Construction | Antibiotic resistance | Reference |
| --- | --- | --- | --- |
| **Sequence derived strains.** |  |  |  |
| R6 | Unencapsulated laboratory strain origin of *blpHC<sub>R6</sub>* |  | 39,40 |
| D39 | Encapsulated serotype 2 clinical isolate |  | 41 |
| TIGR4 | Serotype 4 clinical isolate origin of *blpHC<sub>T4</sub>* |  | 42 |
| P3 | Serotype 6B BlpCP164 strain origin of *blpHC<sub>P164</sub>* |  | 43 |
| 802 | R6 BlpHC<sub>6A.3</sub> strain, origin of *blpHC<sub>6A</sub>* |  | 17 |
| **Reporter strains: *blpC* deletion strains** |  |  |  |
| 1666 | R6 with *PBIR-luc blp* locus with replacement of *blpA-X* with *blpA* intact, *blpQ* from P3 in BIR | cmR, strR | 25 |
| 2434 | P1666 but *blpH<sub>6A</sub>ΔblpC* | cmR, strR | 25 |
| 3200 | P1666 but *blpH<sub>T4</sub>blpC::spcR* | cmR, strR, specR | This study |
| 3202 | P1666 but *blpH<sub>R6</sub>blpC::spcR* | cmR, strR, specR | This study |
| 3235 | 2434 but *ΔsepM* | cmR, strR, specR | This study |
| 3245 | 2434 but reinserted wildtype *sepM* | cmR, strR, specR | This study |
| 3237 | 3200 but *ΔsepM* | cmR, strR, specR | This study |
| 3247 | 3200 but reinserted wildtype *sepM* | cmR, strR, specR | This study |
| 3239 | 3202 but *ΔsepM* | cmR, strR, specR | This study |
| 3249 | 3202 but reinserted wildtype *sepM* | cmR, strR, specR | This study |
| 3612 | 3382 but *blpC::spcR* also wt *sepM* replaced | cmR, strR, specR | This study |
| 3610 | 3382 but *blpC::spcR ΔsepM* | cmR, strR, specR | This study |
| **Reporter strains intact *blp* signaling, dual *com*, *blp* reporters** |  |  |  |
| 2287 | P1666 but *blpA_6B-fs ΔbgaA::PcomAB-Nluc* reporter | cmR strR | 25 |
| 3382 | 2287 but *blpHC<sub>P164</sub>* (made with 6B DNA) | cmR strR | This study |
| 3328 | 2287 but *blpHC<sub>T4</sub>* | cmR strR | This study |
| 3483 | 2287 but *blpHC<sub>6A.3</sub>* | cmR strR | This study |
| 3341 | 2287 but $\Delta sepM$ | cmR strR | This study |
| 3342 | 2287 but reinserted wildtype *sepM* | cmR strR | This study |
| 3485 | 3483 but $\Delta sepM$ | cmR strR | This study |
| 3486 | 3483 but reinserted wildtype *sepM* | cmR strR | This study |
| 3597 | 3328 but $\Delta sepM$ | cmR strR | This study |
| 3599 | 3328 but reinserted wildtype *sepM* | cmR strR | This study |
| 3387 | 3382 but $\Delta sepM$ | cmR strR | This study |
| 3388 | 3382 but reinserted wildtype *sepM* | cmR strR | This study |
| 3761 | 1666 but *blpHC<sub>T4</sub>* (*blpA* intact, no *com* reporter) | cmR strR | This study |
| **Incubating strains** |  |  |  |
| 1821 | D39 incubation strain $\Delta blpT-X::Janus$ | kanR, strS | 25 |
| 3384 | 1821 but $\Delta sepM$ | kanR, strS | This study |
| 3385 | 1821 but reinserted wildtype *sepM* | kanR, strS | This study |
| **Chimeric strains for peptide secretion** |  |  |  |
| 2420 | R6 but *blp* locus with replacement of *blpA-X* with *blpA* intact, *blpQ* in BIR, $\Delta bgaA::PcomAB-luc$ reporter, *comD_TIGR4* (*comC1D2* chimera) and *blpC::janus*. | cmR, strR, kanR | 25 |
| 2430 | 2420 but *blpH<sub>R6</sub>blpC<sub>6A</sub>* HC-chimera | cmR, strR | 25 |
| 3696 | 2420 but *blpH<sub>R6</sub>blpC<sub>T4</sub>* HC-chimera | cmR, strR | This study |
| 3729 | 2430 but *blpH<sub>R6</sub>blpC<sub>6A-L21X</sub>* early stop codon in *BlpC<sub>6A</sub>* chimera/early stop | cmR, strR | This study |
| Table 4. Strains used in this study. |  |  |  |

### Pre-incubation of peptide with cells

D39*ΔblpT-X::janus* with either wildtype or *ΔsepM* in frame deletion was grown to OD_620_=0.20-0.30 in THY 6.8. One mL of culture was removed and 5 nM of synthetic peptide added to the culture or to THY 6.8 with no cells. The peptide mixture was incubated at 37C for 30 minutes and tubes were centrifuged at 5,000 × g for 10 min. 100 μL of cleared supernatants or peptide in media alone was added to an equal volume of the appropriate *ΔblpC* reporter strain grown in THY 6.8 to OD_620_ = 0.2 pre-grown in media supplemented with catalase and 330 μM firefly luciferin (88294; Thermo Fisher Scientific). The plate was incubated in a Synergy HTX plate reader set to read absorbance at 620 nm and luminescence every 5 minutes at 37C. Values at 20 minutes are shown in Figure 1. For the BlpC_P164_ assay using both wildtype and *ΔsepM* incubating strains, 1000 ng/mL of BlpC_P164_ was used (final concentration 500 ng/mL) and values were read at 1 hour to allow for any digestion by the reporter strain to occur. For *S. mutans* studies, BlpC_P164_, BlpC_R6_ and BlpC_6A_ were all used at a final concentration of 500 ng/mL and values were read at 1 hour post peptide stimulation.

### Peptide stimulation assay and dose-response curve generation

BlpC reporter strains that carry the BIR promoter driving the firefly luciferase gene and engineered to contain either the R6, 6A or TIGR4 version of *blpH* in the *blp* locus in addition to a deletion in the *blpC* gene were used for these assays. These reporter strains will only activate in response to exogenous peptide due to the absence of endogenously produced BlpC. The reporters were grown to OD_620_ ∼ 0.2 in THY supplemented with 330 μM firefly luciferin and catalase. Peptide was pre-added to plates to create either two-fold or five-fold dilutions depending on the reporter and peptide concentration. Plates were read as above and activity at 1 hour post peptide addition was reported. Dose-response curves were generated in Prism 8.0 after normalizing values between 0 and 100%. A Hill slope of 1 was used for a nonlinear fit of log(agonist) vs normalized response. For competition experiments, the BlpH_6A_ or BlpH_R6_ reporters were incubated with a dilution of truncated BlpC as above, but each well had a fixed saturating concentration of full length BlpC of 500 ng/mL. These data were compared with the identical assay but lacking full length BlpC.

### Creation of reporter strains and *ΔsepM* mutant and matched wildtype strains

All PCR reactions for downstream Gibson assembly, transformation, or sequencing applications were performed using Phusion polymerase (NEB, M0530). For PCR reactions in which cells were used as template, the cells were obtained from starter cultures resuspended in sterile water and added to the PCR reaction at a 1:20 (Phusion). For PCR reactions in which crude Gibson assembly product was used as template, the crude Gibson assembly product was added to the PCR reaction at a 1:100 dilution. Gibson assembly was performed using NEB HiFi DNA Assembly master mix (NEB, E2621). Primers were synthesized by IDT. Primers are listed in Table 5.

**Table 5.**
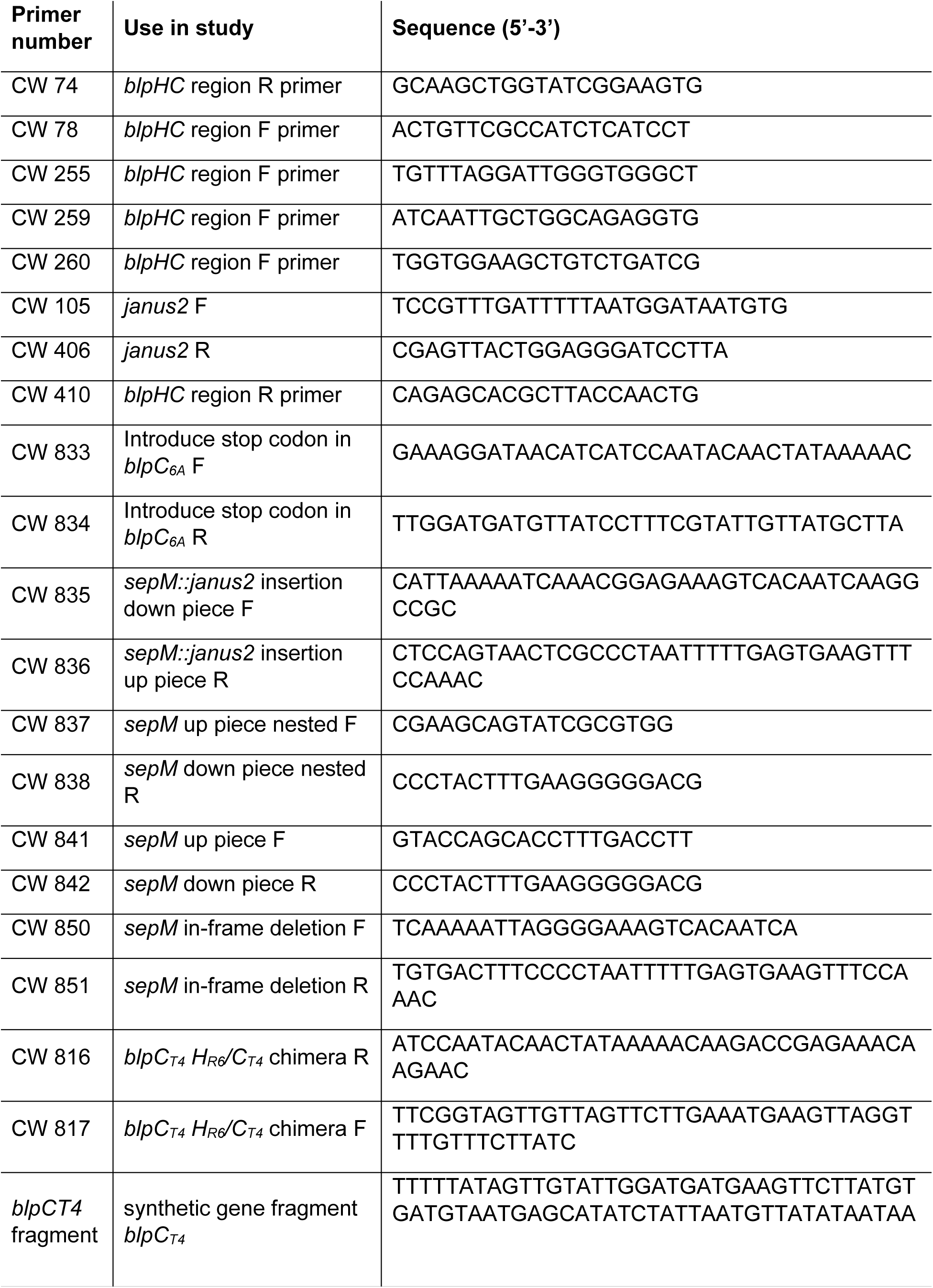
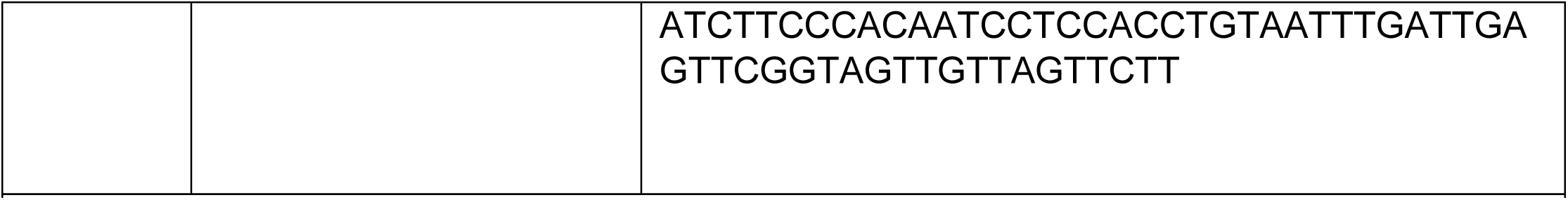
Primers used in this study.

To make *ΔsepM* and matched *sepM*-intact (wildtype) strains, we inserted the Janus2 cassette^37^ between the up and downstream regions of the *sepM* gene using Gibson assembly. The assembled product was reamplified with nested primers and used to transform the relevant strains. The Janus2 cassette is a counter-selectable marker used for allelic exchange that confers gentamicin resistance (200 µg/mL) and sensitivity to 10% sucrose. For subsequent transformation, the Janus2 cassette was moved from otherwise isogenic strains using genomic DNA. An in-frame, unmarked deletion of *sepM* was made using a two-piece Gibson assembly incorporating the deletion and the same nested primers for re-amplification. This product was transformed into the *sepM::*Janus2 strains and selected on sucrose-containing media. To ensure that no unintended mutations occurred during *janus* insertion, the wildtype *sepM* gene was amplified using nested primers and used to transform the *sepM::*Janus2 strains. These strains are referred to as wildtype throughout.

The BlpC_T4_ and BlpC_R6_ Δ*blpC* reporters that link the firefly luciferase gene to the promoter driving the first genes in the BIR region (3200 and 3202 respectively), were made by moving desired sequences into an existing BIR reporter strain with the traditional Janus cassette replacing the *blpHC* region. The Janus cassette confers kanamycin resistance and streptomycin susceptibility^38^. The *blpHC::*Janus strain was transformed with genomic DNA from existing reporters carrying the spectinomycin resistance gene in place of the *blpC* gene as described in Pinchas et al^17^. Strains were selected on spectinomycin and verified by PCR and response to the appropriate BlpC type. To create the *blpC::spectR* BlpH_P164_ strains (3598 and 3600), we used PCR to amplify the spectinomycin cassette in place of *blpC* from an existing *blpH*_*P164*_*blpC::spectR* containing *lacZ* reporter strain^17,25^ with primers CW 410 and 255. We used this PCR product of the *blpHC* region to transform newly created *blpCH*_*P164*_ intact reporters (3387 and 3388) with and without the Δ*sepM* mutation creating strains 3597 and 3599.

To make 3382, 3483, and 3328, the *blp* reporters with intact *blpC* genes used for natural induction assays in the *blpH*_*P164*_, *blpH*_*6A*_ and *blpH*_*T4*_ backgrounds, we moved an existing Janus insertion in the *blpHC* region into the 2287 background. This background has the BIR promoter driving firefly luciferase integrated into the native *blp* locus site and the *comA* promoter driving the nanolux gene integrated into the *bgaA* gene as previously described^25^. This strain background carries the natural frame shift mutation in *blpA*. To make 3382, the Janus was replaced with a PCR product of the *blpHC* region from P3 made using primers CW 74 and 260. Similarly, to make the BlpHC_6A_ and BlpHC_T4_ strains, the region from strains 6A and TIGR4, respectively were amplified with primers CW 259 and 410.

To create the chimeric strains for peptide secretion assays, we used the Janus cassette inserted into *blpC* alone (2420) ^25^ to create the chimeric strains 3696 and 3729 expressing either BlpC_T4_ or BlpC_6A_L21X, respectively. For 3696, we amplified the regions upstream and downstream of *blpC* from the existing BlpH_R6_BlpC_6A_ chimeric strain 2430 using primers CW 78, 410, 816, and 817. We then used a three-piece Gibson assembly to attach it to the synthetic fragment containing the BlpC_T4_ sequence. This piece was used to replace the Janus cassette in *blpC*. For 3729, we used primers CW 78, 410, 833, and 834 to amplify from 2430. Primers CW 833 and 834 introduced a stop codon at the expected SepM cleavage site in *blpC*_*6A*_. We used a two-piece Gibson assembly to fuse the two pieces and transformed that fragment into 2420. ‘The *blpHC* region of both chimeric strains generated in this work were sequenced to confirm incorporation of the desired sequence.

### Natural *blp* and *com* induction assays

To assess for natural *blp* activation (without addition of BlpC peptides), strains were initially grown in THY 6.8 to an OD_620_ of ∼0.2-0.3. 200 μL of this culture was added to 4mL of CDM+ at pH 7.2 with catalase and luciferin added as above. 200 μL aliquots were distributed into a 96 well plate. Luminescence and OD_620_ were read every 5 minutes and plates were incubated at 37C. To assess for *com* activation, nanolux was added to the above media at 1:5000 dilution. Using the Synergy HTX plate reader, the Nluc signal originating from the *comA* promoter driven nanolux gene was isolated using a 450/50-nm band-pass (BP) filter (7082208; BioTek). The RFluc signal originating from the *BIR* promoter driving firefly luciferase was isolated using a 610-nm long-pass (LP) filter (7092209; BioTek) at each reading as previously described^24^.

### BlpC secretion assays

To determine the concentration of BlpC secreted following locus stimulation, chimeric strains with a *blpH*_*R6*_ allele driving either BlpC_6A_, BlpC_6Al21X_ or BlpC_T4_ secretion were grown in THY 6.8 to an OD_620_ of 0.2-0.3 and stimulated with 400 ng/mL of BlpC_R6_ and, in some cases, 250 ng/ml of CSP2. After 1 hour of incubation at 37C, cells were pelleted and supernatants were filter sterilized through a 0.22 μm centrifugal filter (Costar, 816). Samples were used to stimulate *sepM* intact BlpC_6A_ or BlpC_T4_ reporter strains 2434 or 3200. Reporter strains were grown as above in THY 6.8 at 37C to OD_620_ 0.2-0.3 with catalase and luciferin added to media prior to peptide addition. Peptide mixtures were diluted with cells 1:1 and read for OD_620_ and luminescence at 37C over one hour. To determine peptide concentrations, synthetic BlpC_6A_ or BlpC_T4_ was diluted two-fold or five-fold from 1000 ng/ml to create a dose-response curve alongside the supernatant assays. This curve was used to extrapolate secreted BlpC concentrations in the media. Values for samples and controls were obtained 1 hour post peptide addition.

## Acknowledgements

This work was supported by the Ravitz Advancement Award. SD is additionally supported by the Andrew B Briskin Professorship.

## Supplemental Figure 1

**Figure S1.**
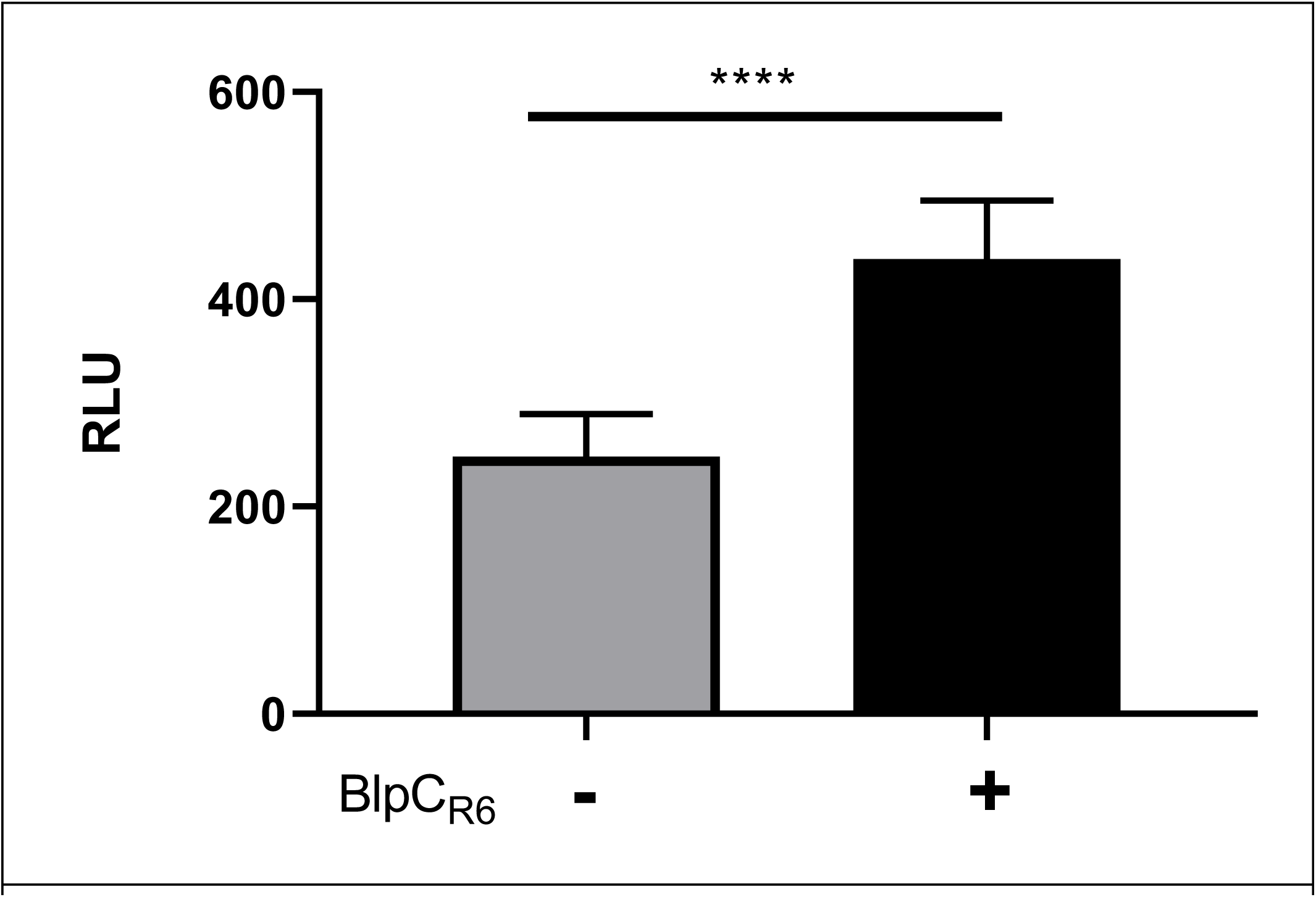
Full length BlpC can activate Δ sepM *blp* reporters at supraphysiologic concentrations to low levels of activity. Activity of *blpH*_R6_Δ*blpCsepM* reporters 1 hour after addition of 1000 ng/mL full length BlpC_R6_ or vehicle control. P value <.0001 by unpaired student T test.

